# A Simulation Study Investigating Potential Diffusion-based MRI Signatures of Microstrokes

**DOI:** 10.1101/2020.08.19.257741

**Authors:** Rafat Damseh, Yuankang Lu, Xuecong Lu, Cong Zhang, Paul J. Marchand, Denis Corbin, Philippe Pouliot, Farida Cheriet, Frederic Lesage

**Affiliations:** Laboratory of Optical and Molecular Imaging, École Polytechnique de Montréal, 2900 Edouard Montpetit Blvd, Montréal (Québec), Canada, H3T 1J4; Montreal Heart Institute, 5000 Rue Bélanger, Montréal (Québec), Canada, H1T 1C8; Université de Montreal, 2900 Edouard Montpetit Blvd, Montréal (Québec), Canada, H3T 1J4; Department of Computer and Software Engineering, École Polytechnique de Montréal, 2900 Edouard Montpetit Blvd, Montréal (Québec), Canada, H3T 1J4

## Abstract

Recent studies suggested that cerebrovascular micro-occlusions, i.e. microstokes, could lead to ischemic tissue infarctions and cognitive deficits. Due to their small size, identifying measurable biomarkers of these microvascular lesions remains a major challenge. This work aims to simulate potential MRI signatures combining arterial spin labeling (ASL) and multi-directional diffusion-weighted imaging (DWI). Driving our hypothesis are recent observations demonstrating a radial reorientation of microvasculature around the micro-infarction locus during recovery in mice. Synthetic capillary beds, randomly- and radially-oriented, and optical coherence tomography (OCT) angiograms, acquired in the barrel cortex of mice (n=5) before and after inducing targeted photothrombosis, were analyzed. Computational vascular graphs combined with a 3D Monte-Carlo simulator were used to characterize the magnetic resonance (MR) response, encompassing the effects of magnetic field perturbations caused by deoxyhemoglobin, and the advection and diffusion of the nuclear spins. We quantified the minimal intravoxel signal loss ratio when applying multiple gradient directions, at varying sequence parameters with and without ASL. With ASL, our results demonstrate a significant difference (p<0.05) between the signal-ratios computed at baseline and 3 weeks after photothrombosis. The statistical power further increased (p<0.005) using angiograms measured at week 4. Without ASL, no reliable signal change was found. We found that higher ratios, and accordingly improved significance, were achieved at lower magnetic field strengths (e.g., B0=3) and shorter readout TE (<16 ms). Our simulations suggest that microstrokes might be characterized through ASL-DWI sequence, providing necessary insights for posterior experimental validations, and ultimately, future translational trials.

## Introduction

Cortical microvascular networks are the carrier of continuous supply of oxygen and energy substrates to neurons, and thus they are responsible for maintaining their healthy state. These networks react dynamically to meet the rapid and substantial increases in energy demands during neuronal activation through the process of neurovascular coupling^1^. Structural deterioration of the cortex microvasculature directly disrupts the regulation of cerebral blood flow and alters the distribution of oxygen and nutrients. Among pathogenic outcomes in cerebrovascular diseases^2^ is the emergence of micro occlusions in penetrating arterioles descending from the pial surface. Recent experiments have provided evidence about the impact of these microscopic events on brain function^3^. Occlusion of a single penetrating vessel was shown to lead to ischemic infarction in the cortex^4^ and to have effects on targeted cognitive tasks. Cerebral microinfarcts have emerged as a potential determinant of cognitive decline, as they are one of the most wide-spread forms of tissue infarction in the aging brain^5^. These cortical lesions have been associated to severe deficits in motor output at muscles^6^. It was also shown that a microembolism of a single cortical arteriole induces cortical spreading depression, a potential trigger and putative cause of migraine with aura^7^. In a separate study, the induction of microvascular lesions in an Alzheimer’s mouse model was shown to alter both the deposition and clearance of amyloid-beta plaques. Optical microscopy and photoacoustic imaging are potential techniques for imaging the local architecture of cerebrovascular morphology at micro-scale, however, they remain invasive and are currently limited to preclinical studies. Given the strong association between these microvascular events and many neurological disorders, developing non-invasive and translatable approaches is of vital importance to identify their presence in clinical settings.

A recent study based on 2-photon microscopy has illustrated that the capillary bed in microvascular networks regenerates into a radially organized structure following a localized photothrombotic infarction^8^. To overcome limitations due to fluorescent dye leakage through the damaged blood-brain barrier, recent work exploiting optical coherence tomography (OCT) provided a detailed exploration of the microvascular angio-architecture rearrangement at different cortical depths^9^ following photo-thrombosis. This latter study confirmed the presence of highly radially organized patterns, at all cortical depths, with a higher degree of structural reorganization in deeper regions. These morphological features could be exploited as clinical signatures of the associated ischemic events.

In this study, we hypothesize that these vascular re-orientations can be detected through magnetic resonance imaging (MRI), and we provide a proof-of-concept through simulations. Our assumption is based on diffusion-weighted imaging (DWI), which is an established MRI technique that provide contrast sensitive to the motion of water molecules^10^. DWI is widely applied to capture white matter tracts^11^, through the use of arbitrarily selected directions of the diffusion gradient to measure the directional bias of molecular movements. A sub-type of DWI is the intra-voxel incoherent motion (IVIM) technique, used to detect the high pseudo-diffusion coefficient that is attributed to the vascular component of the tissue^12^. Pseudo-diffusion measurements were used to quantify changes in cerebral perfusion^13^. Previous studies have shown that a hybrid scheme, combining IVIM and multi-direction DWI, could allow for a measurable effect size caused by microcirculation architecture^14–16^. Phantom-based simulations and realistic measurements of calf muscle were performed to characterize capillary anisotropy of skeletal muscle microvasculature^14^. A further in-vivo study used the same approach to characterize microvascular renal flow anisotropy^15^. Arterial spin labeling (ASL) was combined with an IVIM model to show that such technique is able to capture the dominant directionality of cerebral microvasculature in the rat brain^16^. Here, we conduct IVIM spin echo realistic simulations, to investigate potential signatures of cerebrovascular micro-occlusions induced in mice brains after targeted photothrombosis. Taking advantage of the radial angiogenesis around the micro-infarction locus after occlusion, we propose a measurable biomarker based on quantifying the ratio of directional signal loss induced when using multiple gradient directions. By integrating and excluding ASL, we performed parametric simulations using different field strengths, readout times, b-values and gradient duration. This study provides meaningful insights about the effect size as a function of parameters, which could be of great benefit for guiding further experimental investigations.

## Results

### Differences between diffusion MRI responses due to altered vascular orientations

Exploiting Monte-Carlo MRI simulations, on synthetically approximated capillary beds, we show that orientations of vascular geometry lead to different profiles of the diffusion MRI signal. We constructed two sets (30 samples each), see Figure 3 (B), of synthetic microvessel tubes, with distinct orientional structures. We aimed at mimicking the dissimilarity between healthy and post-stroke angiogenesis occurring before and after the induction of thrombotic occlusions on a penetrating artery within a microvascular unit. The first synthetic set contained randomly oriented vascular segments, which one would expect in healthy microvascular networks. On the other hand, samples in the second set were designed to follow radial orientations observed from realistic post-occlusion OCT acquisitions (see Figure 1 (D)). Random vessel radii ranging from 2-4 *μ*m and arbitrary flow directions were assigned to capillary segments in our synthetic data. We approximated flow and intravascular PO2 as described in the Methods section (see Figure 3 (C)). We extracted an MRI response simulating several gradient directions uniformly distributed in space by varying *θ* 1 and *θ* 2 as depicted in Figure 3 (A). Our simulations show variations in signal behavior as a result of changing gradients associated with different vascular orientations. An example of simulated output from both randomly- and radially-oriented samples is shown in Figure 3 (A). It is clear that after the second diffusion gradient following the 180° pulse, signals recover at different rates depending on gradient directions. When the direction is perpendicular to the vascular flow, advection of spins had no contribution on signal loss. On the other hand, diffusion gradients that had components aligned with flow direction introduced a noticeable loss in signal intensity. This observation suggests that a quantification of the differences between signal readouts at TE time point can uncover useful information about vascular orientation, and thus be used as a signature of angioarchitecture changes associated with microthrombosis.

**Figure 1.**
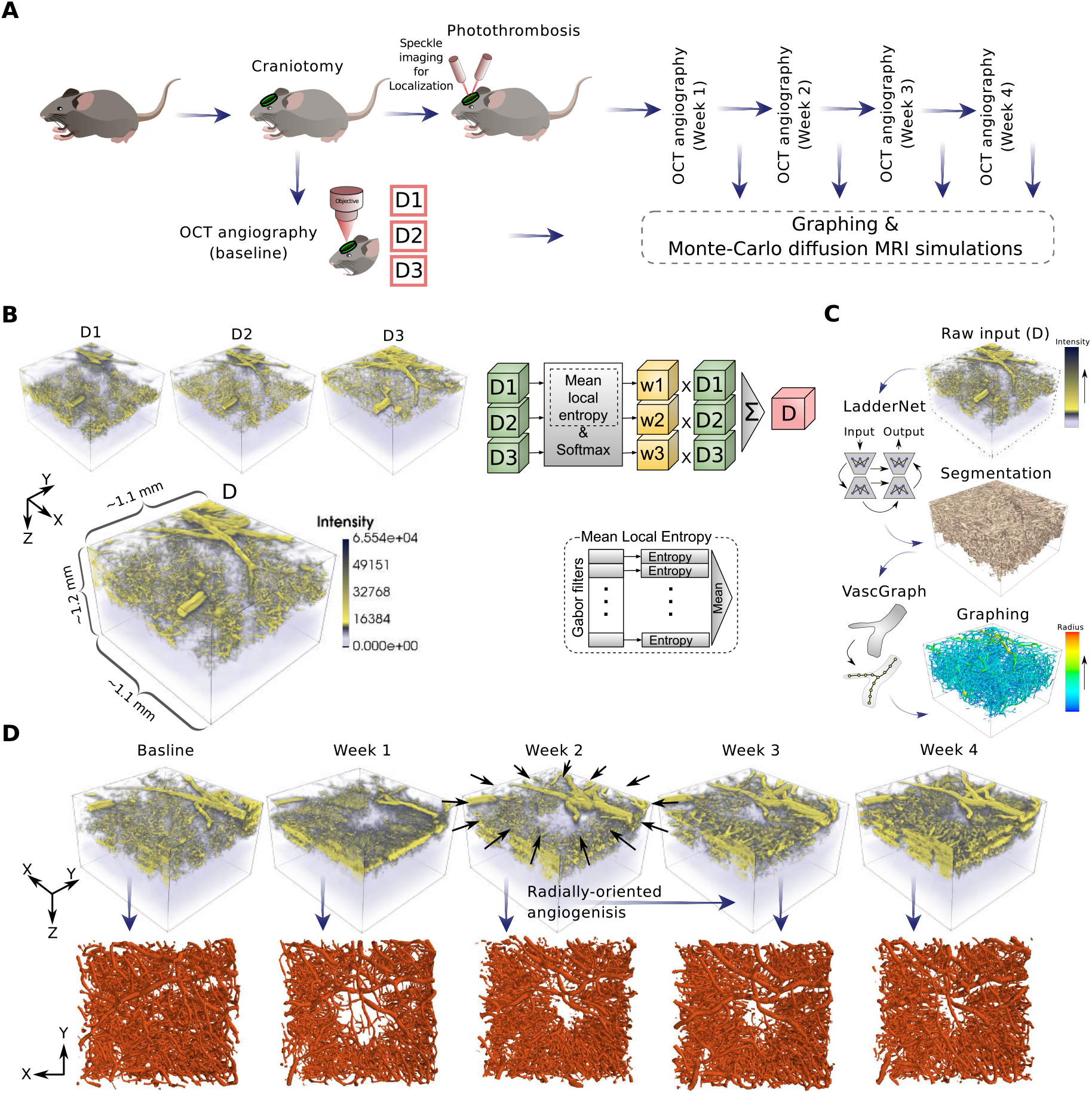
(A) Our experimental procedure for inducing and monitoring of micro-occlusions. Depth-dependant Pre- and post-lesion OCT angiographic acquisitions were preformed to capture vascular degeneration. The OCT stacks acquired at different time points are fed to our computational pipeline to study differences in their diffusion MRI response. (B) Our technique for reconstructing a final 3D OCT angiogram from the three depth-dependent images. We computed their mean local entropy after processing with a set of Gabor filters; a patch with richer vascular structures contributes more to the weighted sum. (C) Our image processing pipeline used to extract useful structural/topological models of vascular networks. These models are essential to perform Monte-Carlo MRI simulations. The Segmentation is based on a customly trained LadderNet architecture. We used The VascGraph toolbox^17^ to obtain graph-based vascular skeletons that can approximate the needed anatomical information. (D) 3D rending of the vascular structure before and after creating a photothrombotic lesion. A noticeable radial-wise orientation is observed after-lesion especially following Week 2.

### Effect size using different MRI parameters

To investigate the statistical significance between the randomly- and radially-oriented synthetic samples based on their diffusion-based MRI responses, we quantified the sample-wise maximum difference of signal loss at readout, *max*(*S*_*i*_/*S*_0_) − *min*(*S*_*i*_/*s*_0_), (see the Methods section and Figure 2 (B)). Larger differences imply more anisotropic orientations (i.e., radially structured in our case). We used a non-parametric Mann-Whitney test to examine the statistical significance between *ϕ* values associated with the two synthetic sets. We retained the set of p-values calculated after using a subspace of B0, TE, *δ* and *b* (see Figure 3 (D)). We repeated the experiment with and without eliminating signal contributions of spins in the extravascular space, i.e., using ASL. As anticipated, we observed larger effect sizes, in general, when using ASL. The selection of the TE value had a key role in increasing the difference between the two set of measurements. The lower the TE value, the greater the statistical significance. A comparable effect was observed of the field strength B0: a higher B0 reduced the difference between the two configurations. Furthermore, an increased effect size was observed when using longer gradient time *δ*. Conversely, the effect associated with the various b-values on effect size was unnoticeable.

**Figure 2.**
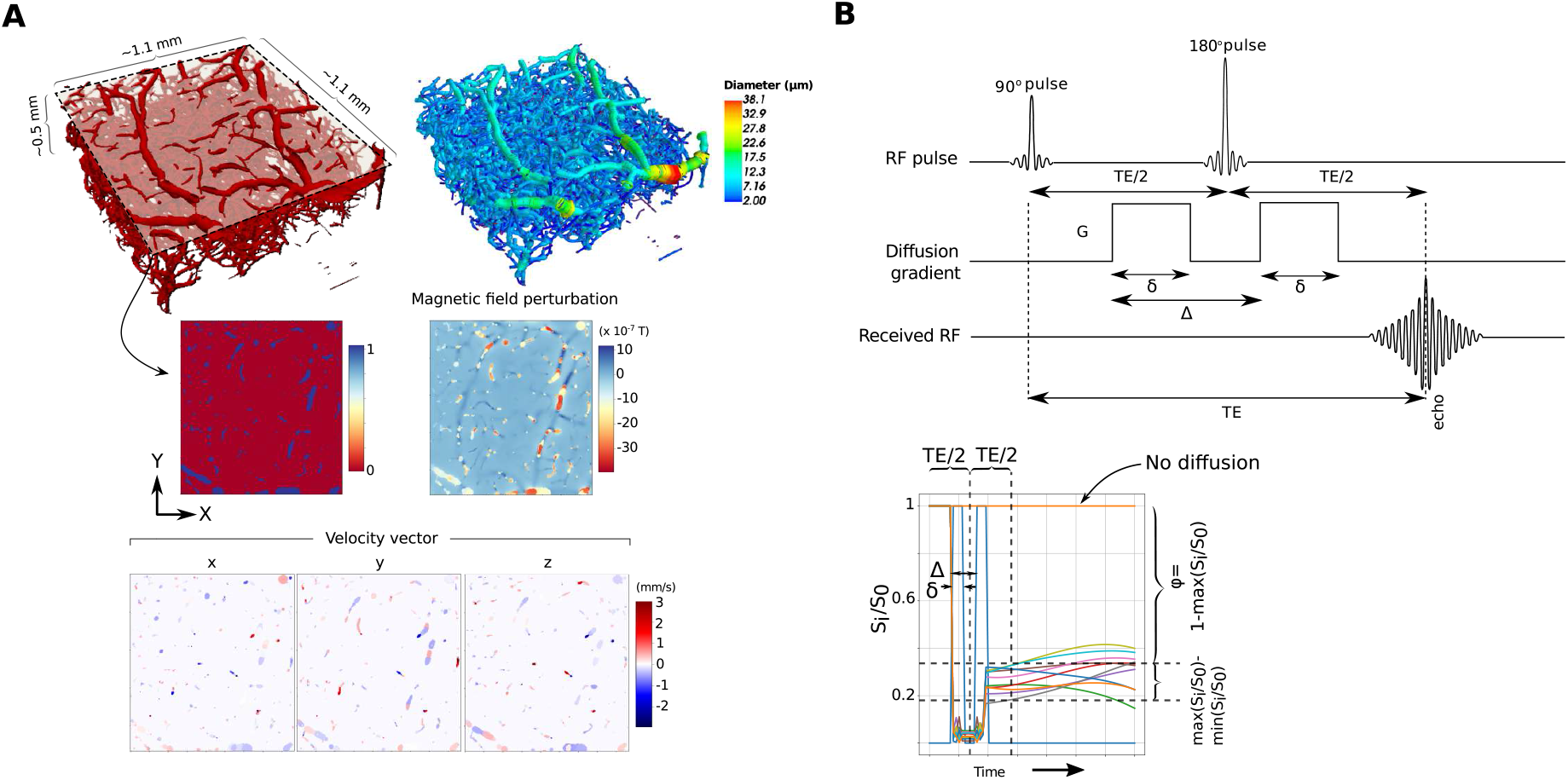
The simulation framework used to compute the MRI response through the diffusion and advection of nuclear spins within the cerebral microvasculature. (A) The computation of alterations in the main magnetic field due to the distribution of deoxyhemoglobin in the blood. These perturbations are calculated based on the PO2 values approximated using our random forest model (see Figure 3). Following the same machine learning approach, we estimated the velocity field that drives the advection of the spins. (B) The quantification of signal variations after simulating the diffusion MRI responses of a vascular unit using different gradient directions; above is the DWI sequence used for each gradient direction.

**Figure 3.**
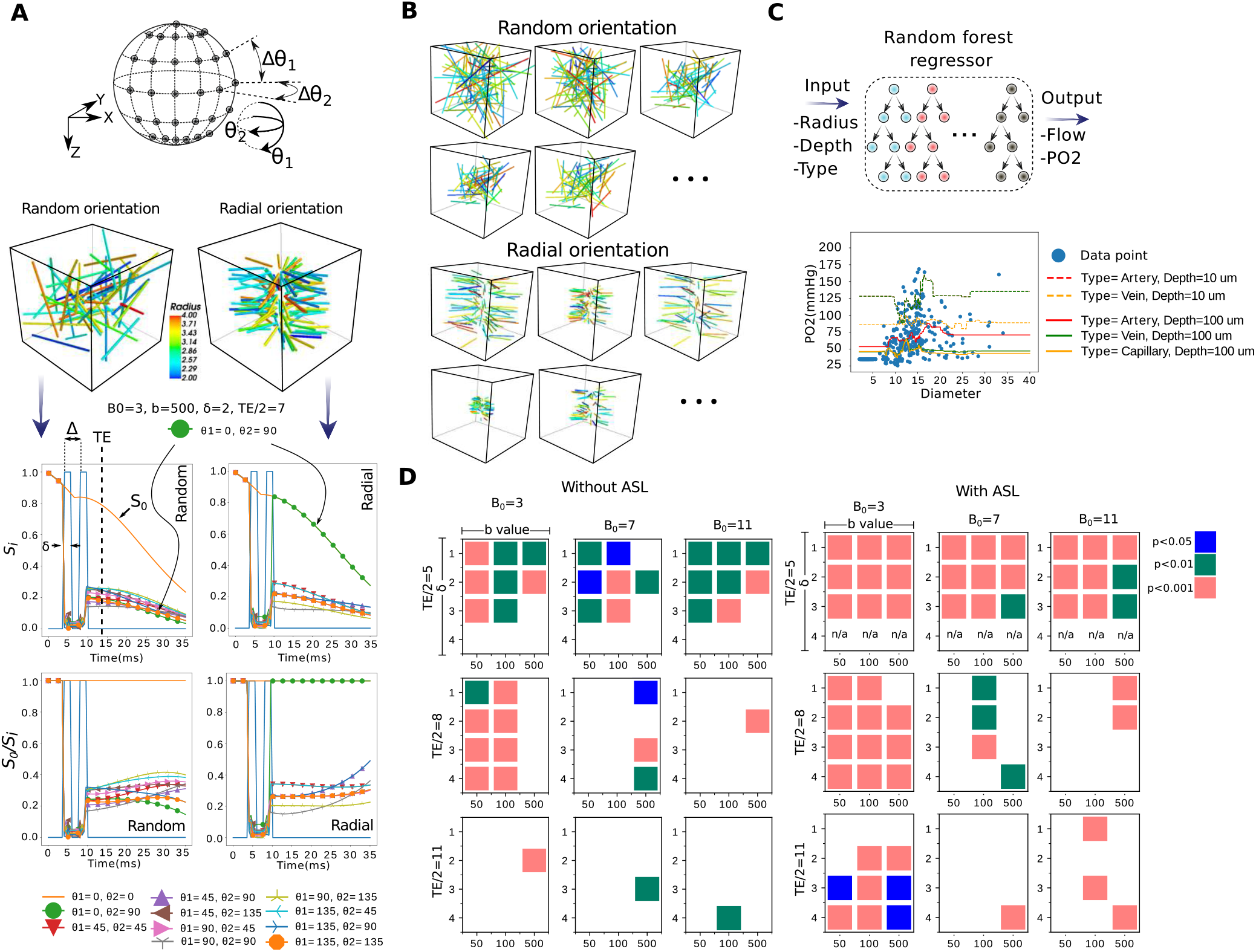
(A) A plot describing the difference between two diffusion-based MRI responses simulated -using a Monte-Carlo framework- from synthetic randomly- and radially-oriented capillary structures. For each sample, we simulate signals resulting from using different gradient directions controlled by the angles *θ* 1 and *θ* 2. (B) Examples from the two groups in our synthetic dataset. (C) A description of the random forest regression model used to approximate Flow and PO2 across the all branches/segments in our vascular models. The model predicts these values for a vascular element based on its annotated radius, depth and vessel type information. (D) Using a subspace of parameters that determine our diffusion MRI sequence, we plot the corresponding statistical p-values computed between our two synthetic groups based on their simulated *ϕ* values.

### Diffusion-based MRI signatures of microstrokes using realistic simulations

Exploiting acquisitions of the angioarchitecture acquired longitudinally following photo-thrombosis, we then investigated the capability of the above sequences at detecting a longitudinal change in microvasculature that reflects in vivo conditions. Here, we report the statistical differences, in terms of the measured *ϕ* values (see Figure 2 (B)), between healthy (baseline) and after-lesion (at week 1, 2, 3 and 4) acquisitions. We performed our experiments with and without involving ASL, using different sequence parameters, by varying the field strength B0 and the b-value while setting TE, *δ* and Δ to 16 ms, 3 ms and 6 ms, respectively. To quantify the values of *ϕ* from each sample, we simulated uniformly distributed gradient directions with *δθ* = 30° (see Figure 3 (A)). From Figure 4, it is observed that no reliable difference was achieved when excluding ASL from our simulations. On the other hand, the use of ASL suggests that the proposed marker, *ϕ*, is effective in differentiating between samples at baseline and after the 3rd and 4th weeks of occlusion. For example, at B0=3 and b=500, *ϕ* = 0.1533±0.0083, whereas it is 0.06978 0.01890 and 0.06148 0.01187 at week 3 and 4, respectively. Noticeably, the scaling of *ϕ* increases with lower B0 field strengths and higher b-values.

**Figure 4.**
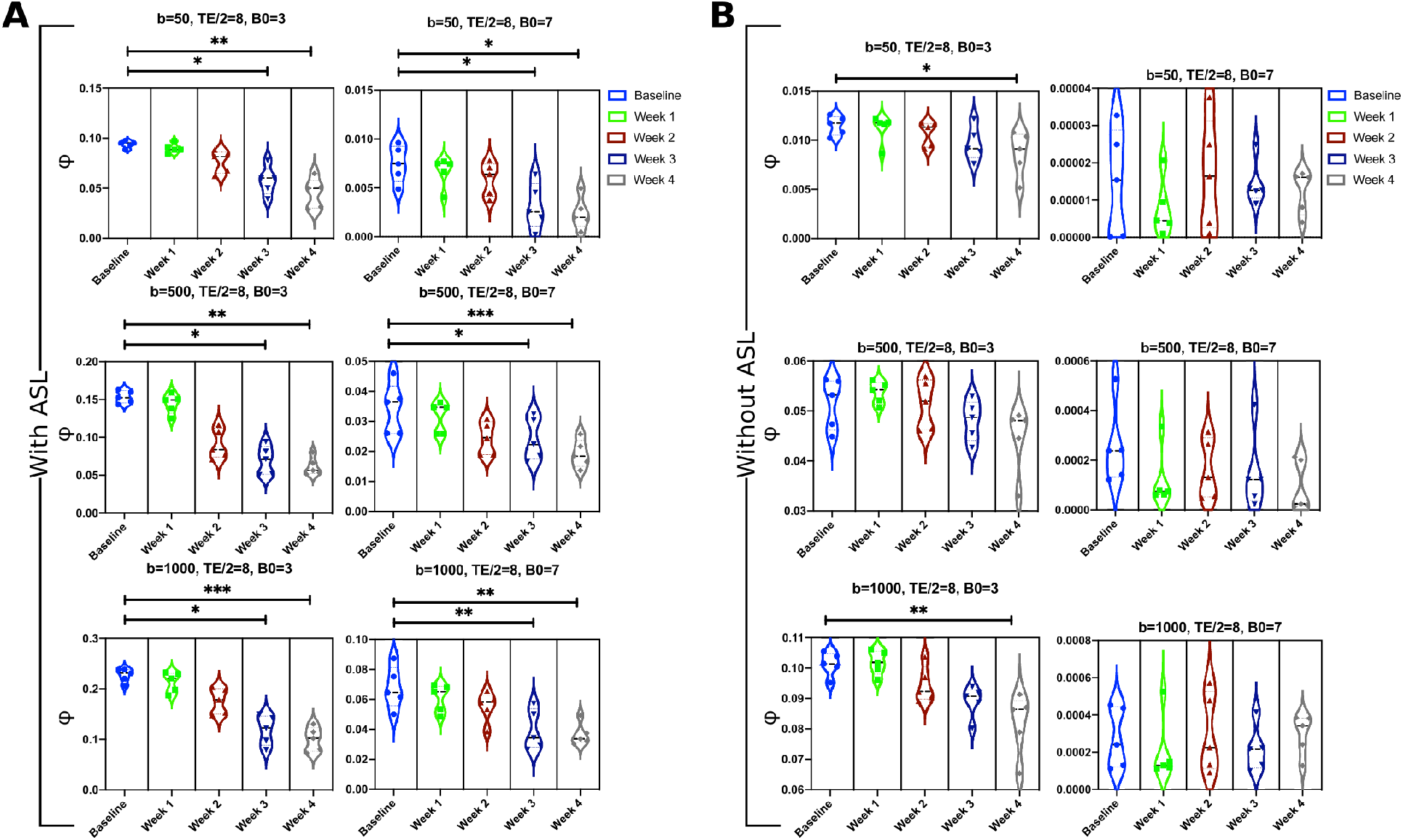
The ratio of minimal signal loss *ϕ* (× 100%) simulated from our OCT angiograms per (or before) and after occlusion through a the multi-directional IVIM scheme (see Figure 2 (B)). We used a set of uniformly distributed gradient directions (Δ*θ* = 30°, see Figure 3 (A)). We propose this measurement as signature distinguishing healthy from lesioned samples. We carried out the Friedman’s test followed by post-hoc comparisons to study the statistical significance between the ratio obtained at the baseline and that calculated at the following 4 weeks after occlusion. Our analysis was performed with and without involving ASL (A) and (B), respectively.

## Discussion

We leveraged OCT microscopic imaging and Monte-Carlo simulations to study potential diffusion MRI signatures of microvascular architecture after experimentally inducing photothrombosis. Our results support the hypothesis that such biomarkers are achievable, under certain assumptions, by exploiting the radial arrangement of microvascular capillary compartments post-lesion. Thus this work suggests that quantifying the differences in signal readouts arising from utilizing multiple gradient directions can characterize such vascular orientations. Simulation outputs provided useful insights about the prospective experimental implications. The use of ASL is of critical importance when characterizing microvascular occlusions based on the associated disruption in vascular geometry. In the mouse cerebral cortex, microvascular density sums to less than 0.05 *mm*^3^ of a tissue volume of 1*mm*; this ratio decreases with depth^18^. Eliminating the MRI signal contributed from the extravascular space is thus necessary. ASL approaches have already shown promising applications in measuring regional cerebral blood flow (rCBF) or perfusion^19–21^, and in assessing vascular disorders^20, 22, 23^. Our simulation outcomes justify the selection of ASL-coupled multi-directional diffusion MRI used by Well et al.^16^ to annotate flow patterns in the mouse cerebral cortex. We observed a noticeable drop in statistical significance between the responses of healthy and lesioned vasculature at longer echo time TE. This is due to the dominant T2/T2* shortening caused by deoxyhemoglobin distribution, opposed to that rising from diffusion gradients. A similar conclusion of reduced statistical differences was obtained when examining the responses at higher B0values. Ultra-high field strengths, despite improving signal-to-noise (SNR) ratio, translate into shorter T2* and T2^24, 25^. In other words, The increased B0 inhomogeneity at higher B0 leads to more signal loss, especially with longer echo times, which offset the advantage of its higher SNR. A straightforward approach would employ shorter echo time to mitigate the adverse effect of stronger fields^26, 27^.

No clear association has been found between the b-value and the statistical difference used to distinguish normal from lesioned samples. More comprehensive investigations are conditioned on further improvements of our simulation framework. We employed a machine learning approach to predict SO2 and blood flow distributions across our vascular networks to enable this study. These measurements can be improved through oxygen transport modeling based on more adequate vascular computational graphs. It is to be mentioned that such modeling remains challenging since it requires tedious manual efforts and can be infeasible at scale^28^. It is known that larger microstrokes impose T2 changes in tissue. The proposed modeling could be improved through the incorporation of measured T2 tissue changes to understand their impact with a more realistic simulation. Another aspect of improvement is related to integrating a model of the restricted diffusion in tissue, instead of assuming a constant extravascular T2 field. An improved framework could encompass the magnetic perturbations induced bysusceptibility interfaces between vessels and cells, and the permeability of the vessel wall^29^.

## Methods

### Animals

All procedures were approved by the Animal Research Ethics Committee of the Montreal Heart Institute. Animal experiments were performed complying with the Canadian Council on Animal Care recommendations. Five C57BL/6J male mice of age 3-6 months were used. Cranial window implantation was carried out for each mouse over its left barrel cortex (0.5mm posterior to bregma, 3.5mm lateral to the midline) to perform OCT imaging. Following scalp retraction, a craniotomy with a diameter of 3 mm was done using a micro-drill and the dura was kept intact. We covered the exposed brain surface with a stacked four-layer glass cover slip (3×3mm, 1×5mm diameter) and sealed it with dental acrylic cement to prevent potential infection. A fixation bar was glued to the skull using the dental acrylic. During surgery, physiological parameters, including electrocardiogram, respiration, heart rate and oxygen saturation of the isoflurane-anesthetized mouse were continuously monitored by a small animal physiological monitoring system (Labeo Technologies Inc. Canada), whose heated platform module also maintained the mouse body temperature at 37 °C. OCT acquisitions were performed on awake resting mice to avoid the modulation of vascular and neural physiology^30, 31^ by anesthetics. During image acquisition, the mice were placed on a free treadmill wheel with their head fixed on a metal frame by the surgically attached bar. It is to be mentioned that OCT-angiography is phase-sensitive and that even sub-pixel motions can dramatically diminish signal to noise ratio (SNR), and hence it is important that the mice stay still during imaging sessions. Accordingly, we trained the mice for head restraint prior to OCT measurements to habituate them to head fixation and reduce their stress. After a week of training on the treadmill wheel, the mice were able to reach a resting state within five minutes after being fixed onto the setup. They were able to stay calm and still for periods of minutes separated by short bouts of locomotion. After the initial baseline measurement, the mice were still trained every day between imaging sessions to maintain their habituation to head restraint throughout the study. The mice were closely monitored for locomotion during image acquisitions.

### Ischemic stroke model

The stroke model exploited a localized photo-thrombosis procedure which is based on a photochemical reaction introduced by Watson et al.^32^. Mice were first intraperitoneally administered Rose Bengal (15 mg/mL, 0.2 ml), a photosensitive dye. A selected cortical region, free of large vessels, was irradiated by a focused green laser beam, since large-vessel thrombosis could lead to a less predictable and less controlled outcome. In addition, avoiding regions with large pial vessels could also minimize the effect of tail artifacts in OCT angiography images^33^. Green light illumination enforces Rose Bengal to produce free radicals that lead to a damage in the endothelium of the microvasculature, thereby triggering discoid platelet aggregation that eventually leads into thrombotic occlusions. The whole process of photo-thrombosis was managed and monitored using a home-built laser speckle imaging system.

### OCT Acquisition system

Imaging of cortical structure and vasculature was performed with a home-build spectral-domain OCT. A broadband light source centered at 1310 nm from a superluminescent diode (SLD) (LS2000C, Thorlabs, USA) was split between the sample arm and the reference arm by a 90:10 fiber optic coupler (TW1300R2A2, Thorlabs, USA). A long working distance objective (M Plan Apo NIR 10X, Mitutoyo, Japan) was installed at the end of the sample arm to focus the collimated light beam into the tissue sample. The spectral interferogram was registered by a spectrometer (Cobra 1300-[1235-1385 nm], Wasatch Photonics, USA) and then digitized by a frame grabber (PCIe-1433, National Instruments, USA). Dispersion mismatch between the two arms was first carefully compensated with N-SF11 compensation glass (Edmund Optics, USA), and the small residual mismatch was then finely corrected with a numerical compensation technique^34^. The axial resolution was measured to be about 4.15 *μ*m in biological tissues. The lateral resolution in tissue was about 2.3 *μ*m. In the sample arm, a dichroic filter was placed to transmit the infrared light used by the OCT system and deflect the visible light for wide-field imaging. The wide-field imaging helped locate the region of interest (ROI) to be scanned by OCT. The sample arm consisted of a galvanometer scanner, a beam expander and an objective lens. The arm was mounted on a motorized vertical translation stage (MLJ150/M, Thorlabs, USA). Adjusting the depth of the imaging focal point can be performed by elevating or lowering the vertical stage. The treadmill wheel onto which the mouse was attached was fixed on a motorized XY linear translation stage (T-LSR, Zaber Technologies, Canada) for fine adjustment of the relative lateral position of the cranial window with respect to the light beam. The 3-axis motion control was integrated into our acquisition software.

### OCT angiographies

We scanned a 1 mm × 1 mm region with the photo-thrombosis-induced lesion located in the center. Our Volumetric scans of the cortex contained 450 B-frames, each of which was composed of 500 A-lines. First, raw spectra were resampled in k-space and then multiplied by a Hanning window. Then, inverse Fourier transform (IFT) was applied to obtain 3D complex-valued OCT structural images. B-scans were repeated twice at each position along the slow axis. Global phase fluctuations (GPF) caused by sub-pixel motion within repeated B-frames were corrected based on the assumption that dynamic tissue only accounts for a very small percentage of brain tissue and that phase and intensity of light reflected from static tissue remain constant^35^. In principle, light reflected by moving red blood cells (RBC) experiences a large phase shift and/or a big intensity change. Thus, we obtain a vascular image by taking the phase and intensity difference between GPF-corrected repeated B-frame^36^. The resulting 3D angiograms were filtered with a 3D Gaussian smoothing kernel with a standard deviation equal to 1 pixel in all three dimensions. A fast axis scan rate of 90 Hz was used, which resulted in an acquisition time of 10 seconds per volumetric scan. In order to extract more comprehensive information of the capillary network in the cortex, three time-resolved OCT-angiographies were performed in the same ROI with light beam being focused at three different cortical depths, namely 250 *μ*m, 400 *μ*m and 550 *μ*m beneath the cortical surface. To achieve a shift of the axial focus in the tissue, we preformed a vertical translation of the objective lens in the sample arm mounted on the vertical translation stage. After, OCT stacks for each animal that were taken considering three different depth-dependent setups, are recombined to form one stack. To obtain the final desired stack, we followed a procedure based on measuring the mean of local entropies computed from each stack after normalizing and convolving it with a set of Gabor filters. Mean entropy measures for all the stacks are then normalized with a softmax function imposed on the axis representing the index of each stack. Voxel intensities in each stack are then weighted by the corresponding normalized local entropies. Finally, we take the sum of the weighted intensities from all the stacks to reconstruct the output stack. In our procedure, we used 18 two-dimensional Gabor filters built with orientations ∈ 0, *π*/6, *π*/3, *π*/2, 2*π*/3, *π*, phase offset ∈ 2.5, 5, 7.5 and a wavelength = 0.01. The kernel used for calculating the local entropy is of size (15, 15). We ran slice-wise calculations to quantify the entropy of each stack. In our chronic microstroke study, we performed 6 OCT imaging sessions over 28 days. The baseline measurement was taken one day before photo-thrombosis. The second imaging session took place 3 days after the thrombotic lesion was induced in the mouse brain, and the following 4 measurements were made 8 days, 14 days, 21 days and 28 days respectively after the ischemic stroke event.

### Vascular Segmentation and Graphing

Many works on vessel segmentation have been presented in the literature. The best recent schemes were based on U-Net neural networks^37^ with their convolutional architecture that accepts images of arbitrary sizes. In this work, we used the LadderNet architecture employed in^38^, which is inspired from the U-Net one but with more interconnected information paths. This architecture can be seen as multiple stacked U-Nets with more pathways. Compared to the conventional U-Net scheme, the shared-weights structure in the LadderNet allows horizontal propagation through the stacked U-Nets which forces the learning process to be made in earlier layers, and thus provides better results^39^. We trained and evaluated our network on an in-house prepared and annotated dataset from Two-photon microscopy angiograms. Our dataset consisted of 59 8-bits 2D grayscale images of 256×256 pixels which were then split into 70%, 15% and 15% portions for training, validation and testing, respectively. Images were standardized by subtracting the mean and dividing by the standard deviation. Contrast limited adaptive histogram equalization was then applied to correct light imbalance. Images were adjusted with a gamma value of 1.2. Training, testing and validation processes were done patch-wise (patch size of 32×32) after augmentation with random rotations. Each of the down and up streams paths in the network had 4 convolutional layers. A leaning rate of 0.001 was used. We applied the trained network on each slice in our 3D OCT angiograms to obtain the corresponding delineated vascular structures. After segmentation, we employed a graphing method, proposed recently by Damseh et al.^17^, to transform our binarized inputs into a fully-connected graph-based skeletons. The method initiates geometric grid graphs encapsulated within vascular boundaries and iteratively deforms them toward vessel centerlines. Converged geometries are then refined and converted into graph-based skeletons. The output graph models consisted of nodes distributed along vessel centerlines to capture the geometry and edges connecting these nodes to represent the inherited topology. The method assigns vessel radii to graph nodes with unique identifiers to each set of them located at a certain vascular branch/segment. We used the VascGraph toolbox associated with that work, available on https://github.com/Damseh/VascularGraph, to generate and visualize the anatomical graphs.

### Flow and PO2 Regression

To prepare the generated anatomical model to undergo MRI simulation experiments, we had to assign biophysical quantities across vascular compartments, namely flow and PO2 values, necessary for reconstructing the T2* field. Resolving these values through biophysical simulations is tedious due to the low-quality nature of OCT inputs, which hinders the formulation of a correct geometric domain suitable for such computations. In other words, such computations are extremely sensitive to topological errors that can easily disrupt the final solution. Here, instead, we followed a machine-learning based approach that utilizes an ensemble of random decision trees, i.e, random forests, fitted on experimental data collected from mice models in a previous work performed by Moeini et al.^40^. The data consisted of > 300 measurements of PO2 and Flow values of cerebral microvascular segments having different sizes, types and located at different cortical depths. We built two separate random forest models that accept radius, type and depth information for a vascular segment and output the corresponding flow and PO2 values. Types of vessel segments were determined after thresholding on the radius value of 3 *μ*m^40^. Vascular segments with radii above the threshold were randomly set to be either arteries or veins, whereas the rest were set as capillaries. Following this procedure, we only needed an approximation of the vascular geometry to capture the anatomical features required by the random forest model, and avoided the dependency on the biophysical modeling that requires accurate annotations and is highly prone to any topological error induced in the vascular model. Each of the two random forests consisted of 100 decision trees with 10 maximum depth. The architecture of our regression forests was selected after assessing the mean square error (MSE) resulting from using different architectural setups. The same measure of MSE was used to determine the quality of a split in the single decision trees that compose the forest model. The minimum number of samples required to split in each decision tree has been set to 2. Bootstrapping was used to reduce the variance resulting from the outputs of multiple trees fitted on random sub-samples, with replacement, from the original training set^41^.

### MRI simulations

We implemented a Monte-Carlo MRI simulation code in Python following the same routines of previous protocols^28, 42^, which was built using Matlab. Simulation principles in these works were originally based on descriptions detailed in^43^. The previous implementations were performed to tackle the Blood-oxygen-level-dependent (BOLD) MRI response. Here, to align with our study goals, we developed new functionalities to create multi-directional spin echo DW sequence simulations. In our numerical MRI trails (coinciding with the aforementioned studies), it was assumed that the MRI response in the mouse brain could be approximated by probing the behavior of a large number of hydrogen water protons moving in background due to free diffusion and constrained advection processes. Practically, our assumption incorporated the following in-detail descriptions: the domain enjoys a spatially constant T1 relaxation; T2 value in tissue regions is fixed but varies within the vascular space; T2* effect has spatial variations subjected to the level of deoxyhemoglobin within the vasculature (see Figure 2 (A)). As suggested in previous works designed for studying vascular-based MRI responses^28, 42, 44, 45^, we neglected a set of factors that have insignificant effect on the extracted signal: hydrogen atoms not bound to water, additional iron presence in basal ganglia and some other brain parts, external B0 and B1 fields inhomogeneities, gradient nonlinearity, hematocrit variability. Also, to enable our analysis, we excluded the macroscopic magnetic field destruction occurring due to imperfect shimming or the microscopic field differences emerging from the applied gradients. In our Monte-Carlo simulations, we measured the voxel-wise response based on summing the accumulated complex phase from 1 × 10^−6^ spins, after initiating their positions at *t* = 0 using a uniform probability distribution that covers the spatial domain, and then updating their location every 0.05 ms. The protons were let to diffuse isotropically with a diffusivity constant of 0.8 *μ*m^2^/ms^46^. With the assumptions mentioned above, among the relaxation constants necessary for our computations is that attributed to the local magnetic field inhomogeneities, affecting T2*, due to the presence of deoxyhemoglobin in the vasculature. We followed the numerical method described in^47^ to compute the perturbations in the magnetic field by convolving the susceptibility shift volume, which is calculated depending on the level of PO2, with an ellipsoidal kernel oriented with the field B0. Another important relaxation factor affecting T2 is that related to the spin-spin coupling, where it is computed based on the fitted models illustrated in^48^. In the extravascular space, T2 is fixed and obtained in seconds as

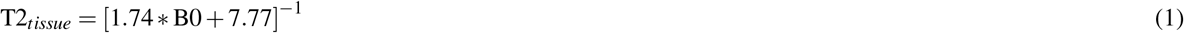

On the other hand, this parameter spatially varies in the vasculature depending on the oxygen saturation level (SO2):

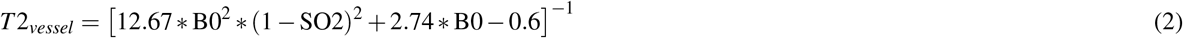

The SO2 values were calculated from their PO2 counterparts based on the Hill conversion equation with coefficients specific for C57BL/6 mice (h=2.59 and P50=40.2)^49^. The relaxation effect due to T1 was included as an exponential decay with T1 =1590 ms^50^. Hematocrit values necessary to compute the field susceptibility as described in^47^ were assumed to be 0.44 in arterioles and venules, and 0.33 in capillaries^51^. Based on the mappings of anatomical features and biophysical quantities computed from OCT angiographies, we computed their MRI response following a combined design of IVIM and multi-direction DWI sequence. The simulated MRI signal is constructed from the contributions of intravascular (IV) and extravascular (EV) spins. Assuming a negligible exchange between the IV and EV components during the echo time^16^, the total DW voxel signal captured at a gradient *i*, with its direction **u**_*i*_ ∈ **U**= {**u**_1_, **u**_2_, …**u**_*n*_}, could be expressed as:

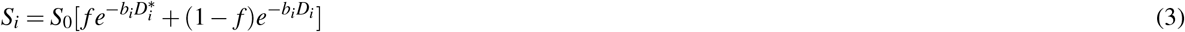

where *S*_0_ is the response at zero diffusion gradient, *f* is the fraction of vasculature in the voxel, and *b*_*i*_(mm^2^/s) is a parameter that characterizes the diffusion gradient *i*. The term *D_i_* is the apparent diffusion coefficient (ADC) influenced by the Gaussian diffusion in tissue, whereas 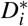 is the pseudo-diffusion coefficient (PDC) representing the perfusion of spins in the microvascular network. A diffusion gradient consists of two short pulses of duration *δ*, with amplitude *G*, separated by a short amount of time Δ. The corresponding b-value can be obtained as:

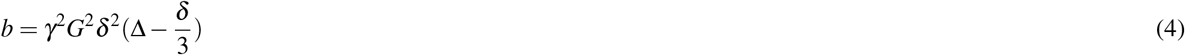

where *γ* is the proton gyromagnetic ratio of Hydrogen. Reflecting this on our task of finding a signature about the deterioration in the microvascular architecture occurring due ischemic thrombotic events, our interest is the set of ratios 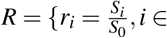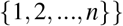, calculated through applying *n* gradients with directions equally sampled from all possible choices in the 3D space. As discussed in the Introduction section, previous works^8, 9^ reported that voxels with thrombotic ischemia exhibit a highly radial orientation of microvessels in the lateral plane around the lesion core. This implies that a higher perfusion rate, i.e., an increased loss in MRI signal, would be observed when a gradient field is parallel to the lateral plane compared to that occurring when it is perpendicular to it. In general, healthy voxels beneath the pial surface are assumed to be composed of randomly oriented capillary segments, and thus would result in comparable MRI signal loss regardless to the direction of the selected gradient. Hence, the ratio *ϕ* = 1 − *max*(*R*) could be regarded as a biological marker distinguishing healthy voxels from those having a thrombotic lesion (see Figure 2 (B)). Beside computing *ϕ* through probing spins behavior in an entire voxel or ROI, we also studied the response when neglecting the extravascular spins, thus omitting the diffusion term in (3). Such procedure could be practically translated with ASL. Based on the mappings of anatomical features and biophysical quantities computed from OCT angiographies, we computed their MRI response following a combined design of IVIM and multi-direction DWI sequence. Simulated MRI signal is constructed from the contributions of intravascular (IV) and extravascular (EV) spins. Assuming a negligible exchange between the IV and EV components during the echo time.

## Acknowledgements

This study was funded by a Canadian Institutes of Health Research (CIHR, 299166) operating grant and a Natural Sciences and Engineering Research Council of Canada (NSERC, 239876-2011) discovery grant to F. Lesage.

## Author contributions statement

R.D. and F.L devised the main conceptual ideas and proof outline. R.D. wrote the manuscript and developed the codes for all the computational blocks in the study, from vascular graphing to MRI simulations, and analyzed the data. Y.L. built the OCT system and performed acquisitions. X.L and Y. L. and C. Z. conducted the craniotomy, the cover glass implantation, and the induction of photo-thrombotic lesions. P. M. did the post-processing of the OCT acquisitions. D. C. prepared the LadderNet segmentation code. All authors reviewed the manuscript.

## Additional information

Correspondence and requests for materials should be addressed to R.D.

## References

1. Hillman, E. M. Coupling mechanism and significance of the bold signal: a status report. Annu. review neuroscience 37, 161–181 (2014).

2. Pantoni, L. Cerebral small vessel disease: from pathogenesis and clinical characteristics to therapeutic challenges. The Lancet Neurol. 9, 689–701 (2010).

3. Summers, P. M. et al. Functional deficits induced by cortical microinfarcts. J. Cereb. Blood Flow & Metab. 37, 3599–3614 (2017).

4. Shih, A. Y. et al. The smallest stroke: occlusion of one penetrating vessel leads to infarction and a cognitive deficit. Nat. neuroscience 16, 55–63 (2013).

5. Taylor, Z. J. et al. Microvascular basis for growth of small infarcts following occlusion of single penetrating arterioles in mouse cortex. J. Cereb. Blood Flow & Metab. 36, 1357–1373 (2016).

6. Anenberg, E. et al. Ministrokes in channelrhodopsin-2 transgenic mice reveal widespread deficits in motor output despite maintenance of cortical neuronal excitability. J. Neurosci. 34, 1094–1104 (2014).

7. Dönmez-Demir, B. et al. Microembolism of single cortical arterioles can induce spreading depression and ischemic injury; a potential trigger for migraine and related mri lesions. Brain research 1679, 84–90 (2018).

8. Schrandt, C. J., Kazmi, S. S., Jones, T. A. & Dunn, A. K. Chronic monitoring of vascular progression after ischemic stroke using multiexposure speckle imaging and two-photon fluorescence microscopy. J. Cereb. Blood Flow & Metab. 35, 933–942 (2015).

9. Lu, Y., Lu, X., Zhang, C., Marchand, P.-J. & Lesage, F. Longitudinal optical coherence tomography imaging of tissue repair and microvasculature regeneration and function after targeted cerebral ischemia. J. Biomed. Opt. 25, 046002 (2020).

10. Jones, D. K. Diffusion mri (Oxford University Press, 2010).

11. Alexander, A. L., Lee, J. E., Lazar, M. & Field, A. S. Diffusion tensor imaging of the brain. Neurotherapeutics 4, 316–329 (2007).

12. Le Bihan, D. Intravoxel incoherent motion perfusion mr imaging: a wake-up call. Radiology 249, 748–752 (2008).

13. Federau, C. et al. Quantitative measurement of brain perfusion with intravoxel incoherent motion mr imaging. Radiology 265, 874–881 (2012).

14. Karampinos, D. C., King, K. F., Sutton, B. P. & Georgiadis, J. G. Intravoxel partially coherent motion technique: characterization of the anisotropy of skeletal muscle microvasculature. J. Magn. Reson. Imaging: An Off. J. Int. Soc. for Magn. Reson. Medicine 31, 942–953 (2010).

15. Notohamiprodjo, M. et al. Combined intravoxel incoherent motion and diffusion tensor imaging of renal diffusion and flow anisotropy. Magn. resonance medicine 73, 1526–1532 (2015).

16. Wells, J., Thomas, D., Saga, T., Kershaw, J. & Aoki, I. Mri of cerebral micro-vascular flow patterns: A multi-direction diffusion-weighted asl approach. J. Cereb. Blood Flow & Metab. 37, 2076–2083 (2017).

17. Damseh, R., Delafontaine-Martel, P., Pouliot, P., Cheriet, F. & Lesage, F. Laplacian flow dynamics on geometric graphs for anatomical modeling of cerebrovascular networks. arXiv preprint arXiv:1912.10003 (2019).

18. Tsai, P. S. et al. Correlations of neuronal and microvascular densities in murine cortex revealed by direct counting and colocalization of nuclei and vessels. J. Neurosci. 29, 14553–14570 (2009).

19. Wang, Y. et al. Regional reproducibility of pulsed arterial spin labeling perfusion imaging at 3t. Neuroimage 54, 1188–1195 (2011).

20. Telischak, N. A., Detre, J. A. & Zaharchuk, G. Arterial spin labeling mri: clinical applications in the brain. J. Magn. Reson. Imaging 41, 1165–1180 (2015).

21. Su, P. et al. Multiparametric estimation of brain hemodynamics with mr fingerprinting asl. Magn. resonance medicine 78, 1812–1823 (2017).

22. Zaharchuk, G. et al. Arterial spin-labeling mri can identify the presence and intensity of collateral perfusion in patients with moyamoya disease. Stroke 42, 2485–2491 (2011).

23. Hendrikse, J., Petersen, E. T. & Golay, X. Vascular disorders: insights from arterial spin labeling. Neuroimaging Clin. 22, 259–269 (2012).

24. Choi, S. et al. Dti at 7 and 3 t: systematic comparison of snr and its influence on quantitative metrics. Magn. resonance imaging 29, 739–751 (2011).

25. von Morze, C. et al. Reduced field-of-view diffusion-weighted imaging of the brain at 7 t. Magn. resonance imaging 28, 1541–1545 (2010).

26. Zhan, L. et al. Magnetic resonance field strength effects on diffusion measures and brain connectivity networks. Brain connectivity 3, 72–86 (2013).

27. Gallichan, D. Diffusion mri of the human brain at ultra-high field (uhf): A review. NeuroImage 168, 172–180 (2018).

28. Gagnon, L. et al. Quantifying the microvascular origin of bold-fmri from first principles with two-photon microscopy and an oxygen-sensitive nanoprobe. J. Neurosci. 35, 3663–3675 (2015).

29. Pannetier, N. A., Debacker, C. S., Mauconduit, F., Christen, T. & Barbier, E. L. A simulation tool for dynamic contrast enhanced mri. PloS one 8, e57636 (2013).

30. Janssen, B. J. et al. Effects of anesthetics on systemic hemodynamics in mice. Am. J. Physiol. Circ. Physiol. 287, H1618–H1624 (2004).

31. Masamoto, K. & Kanno, I. Anesthesia and the quantitative evaluation of neurovascular coupling. J. Cereb. Blood Flow & Metab. 32, 1233–1247 (2012).

32. Watson, B. D., Dietrich, W. D., Busto, R., Wachtel, M. S. & Ginsberg, M. D. Induction of reproducible brain infarction by photochemically initiated thrombosis. Annals Neurol. Off. J. Am. Neurol. Assoc. Child Neurol. Soc. 17, 497–504 (1985).

33. Baran, U., Choi, W. J., Li, Y. & Wang, R. K. Tail artifact removal in oct angiography images of rodent cortex. J. biophotonics 10, 1421–1429 (2017).

34. Szkulmowski, M., Tamborski, S. & Wojtkowski, M. Spectrometer calibration for spectroscopic fourier domain optical coherence tomography. Biomed. optics express 7, 5042–5054 (2016).

35. Lee, J., Srinivasan, V., Radhakrishnan, H. & Boas, D. A. Motion correction for phase-resolved dynamic optical coherence tomography imaging of rodent cerebral cortex. Opt. express 19, 21258–21270 (2011).

36. Zhang, A., Zhang, Q., Chen, C.-L. & Wang, R. K. Methods and algorithms for optical coherence tomography-based angiography: a review and comparison. J. biomedical optics 20, 100901 (2015).

37. Ronneberger, O., Fischer, P. & Brox, T. U-net: Convolutional networks for biomedical image segmentation. In International Conference on Medical image computing and computer-assisted intervention, 234–241 (Springer, 2015).

38. Ronneberger, O., Fischer, P. & Brox, T. U-net: Convolutional networks for biomedical image segmentation (2015). 1505.04597.

39. Zhuang, J. Laddernet: Multi-path networks based on u-net for medical image segmentation (2018). 1810.07810.

40. Moeini, M. et al. Compromised microvascular oxygen delivery increases brain tissue vulnerability with age. Sci. reports 8, 1–17 (2018).

41. Louppe, G. Understanding random forests: From theory to practice. arXiv preprint arXiv:1407.7502 (2014).

42. Pouliot, P. et al. Magnetic resonance fingerprinting based on realistic vasculature in mice. Neuroimage 149, 436–445 (2017).

43. He, X. & Yablonskiy, D. A. Quantitative bold: mapping of human cerebral deoxygenated blood volume and oxygen extraction fraction: default state. Magn. Reson. Medicine: An Off. J. Int. Soc. for Magn. Reson. Medicine 57, 115–126 (2007).

44. Yao, B. et al. Susceptibility contrast in high field mri of human brain as a function of tissue iron content. Neuroimage 44, 1259–1266 (2009).

45. Desjardins, M., Berti, R., Lefebvre, J., Dubeau, S. & Lesage, F. Aging-related differences in cerebral capillary blood flow in anesthetized rats. Neurobiol. aging 35, 1947–1955 (2014).

46. Le Bihan, D. Apparent diffusion coefficient and beyond: what diffusion mr imaging can tell us about tissue structure (2013).

47. Marques, J. & Bowtell, R. Application of a fourier-based method for rapid calculation of field inhomogeneity due to spatial variation of magnetic susceptibility. Concepts Magn. Reson. Part B: Magn. Reson. Eng. An Educ. J. 25, 65–78 (2005).

48. Uludağ, K., Müller-Bierl, B. & Uğurbil, K. An integrative model for neuronal activity-induced signal changes for gradient and spin echo functional imaging. Neuroimage 48, 150–165 (2009).

49. Uchida, K., Reilly, M. P. & Asakura, T. Molecular stability and function of mouse hemoglobins. Zool. Sci. 15, 703–706 (1998).

50. Chugh, B. P. et al. Robust method for 3d arterial spin labeling in mice. Magn. resonance medicine 68, 98–106 (2012).

51. Griffeth, V. E. & Buxton, R. B. A theoretical framework for estimating cerebral oxygen metabolism changes using the calibrated-bold method: modeling the effects of blood volume distribution, hematocrit, oxygen extraction fraction, and tissue signal properties on the bold signal. Neuroimage 58, 198–212 (2011).

